# Circadian regulation of Ca_V_1.2 expression by RORα in the mouse heart

**DOI:** 10.1101/2024.01.15.575657

**Authors:** Estelle Personnic, Garance Gerard, Corinne Poilbout, Anton M. Jetten, Ana Maria Gómez, Jean-Pierre Benitah, Romain Perrier

## Abstract

**Background:** In addition to show autonomous beating rhythmicity, the physiological functions of the heart present daily periodic oscillations. Notably the ventricular repolarization itself varies throughout the circadian cycle which was mainly related to the periodic expression of K^+^ channels. However, the involvement of the L-type Ca^2+^ channel (Ca_V_1.2 encoded by *Cacna1c* gene) in these circadian variations remains elusive.

**Methods:** We used a transgenic mouse model (PCa-luc) that expresses the luciferase reporter under the control of the cardiac *Cacna1c* promoter and analyzed promoter activity by bioluminescent imaging, qPCR, immunoblot, Chromatin immunoprecipitation assay (ChIP) and Ca_V_1.2 activity.

**Results:** Under normal 12:12h light-dark cycle, we observed *in vivo* a biphasic diurnal variation of promoter activities peaking at 9 and 19.5 Zeitgeber time (ZT). This was associated with a periodicity of *Cacna1c* mRNA levels preceding 24-h oscillations of Ca_V_1.2 protein levels in ventricle (with a 1.5 h phase shift) but not in atrial heart tissues. The periodicity of promoter activities and Ca_V_1.2 proteins, which correlated with biphasic oscillations of L-type Ca^2+^ current conductance, persisted in isolated ventricular cardiomyocytes from PCa-Luc mice over the course of the 24-h cycle, suggesting an endogenous cardiac circadian regulation. Comparison of 24-h temporal patterns of clock gene expressions in ventricles and atrial tissues of the same mice revealed conserved circadian oscillations of the core clock genes except for the retinoid-related orphan receptor α gene (RORα), which remained constant throughout the course of a day in atrial tissues. *In vitro* we found that RORα is recruited to two specific regions on the *Cacna1c* promoter and that incubation with specific RORα inhibitor disrupted 24-h oscillations of ventricular promoter activities and Ca_V_1.2 protein levels. Similar results were observed for pore forming subunits of the K^+^ transient outward currents, K_V_4.2 and K_V_4.3.

**Conclusions:** These findings raise the possibility that the RORα-dependent rhythmic regulation of cardiac Ca_V_1.2 and K_V_4.2/4.3 throughout the daily cycle may play an important role in physiopathology of heart function.

## Introduction

The biological rhythmicity is fundamental to almost all organisms on earth and plays a key role in health and disease. With its regular beats determined by intrinsic electrophysiological properties of the cardiomyocytes, the heart is the main physiological metronome that we are aware of. Moreover, periodical oscillations of the cardiac contractility, metabolism and electrophysiology are observed throughout the daily cycle to ensures the heart to adapt to geophysical cues, physical exertion and food intake fluctuations by adjusting both its electrical activities and mechanical function^1^. Notably, it is long-recognized that the cardiac repolarization, independently of heart rate variation, show marked circadian periodicity that predispose to circadian occurrence of ventricular arrhythmias^2–7^. Even if sympathovagal tone, as well as others neurohormonal variations, have been considered as major determinants of these circadian patterns, an innate cellular clock mechanism might also be involved. Indeed, these circadian periodicities are maintained in isolated hearts, tissues and even isolated cardiomyocytes^8–10^. All living organisms from prokaryote to eukaryote have developed within cells throughout the body a highly temporal coordinated integrated network of biological clock to temporal coordination and plasticity of internal biological processes. These biological clocks are highly conserved cell-autonomous autoregulatory transcription-translation feedback loops comprising the interlocked activities of a set of basic helix–loop–helix transcriptional factors (TFs^11^). A first loop is composed of heterodimerized activators (Brain and muscle ARNT-like protein-1, BMAL1 and Circadian locomotor output cycles kaput, CLOCK) that bind E-box response element in the regulatory regions to induce the transcription of their own repressors (Period, PER1/2/3 and Cryptochrome, CRY1/2) in a cycle that repeats itself in approximately 24 h. A reinforcing loop that is controlled by the first one and modulates *BMAL1* transcription, is formed by the activator D-site albumin-binding protein, DBP and the repressor Nuclear factor interleukin 3 regulated, NFIL3 also known as E4BP4 that through D-box response elements regulate the expression of *NR1F1* and *NR1D1/2*, encoding retinoid-related orphan receptor α (RORα) and the nuclear receptors REV-ERBα/β, respectively. REV-ERBα and RORα mediate opposing actions to either inhibit or activate *BMAL1* transcription, respectively, via interaction with a ROR response element (RORE). In addition, the exact functioning of this molecular clock core depends on a growing number of intracellular factors, including kinases and phosphatases that regulate the stability of molecular clock proteins, as well as chromatin and chromatin-modifying complexes, all of which contribute to produce a molecular rhythm. In turn, this core molecular clock regulates the periodic expression of nearly 50% of all expressed genes in a tissue-specific manner^12^. The central clock mechanism is the hypothalamic suprachiasmatic nucleus (SCN), which via neuronal, endocrine, and paracrine signals^13^ synchronizes peripheral tissue own molecular clock. The heart expresses core circadian clock genes in a time sensitive manner^14, 15^, controlling the circadian expression of ∼10% cardiac transcriptome and proteome^16–21^. The importance of the cardiac molecular clock has been emphasized by generating cardiac-specific circadian gene knockout animals^6, 22–26^. Remarkably, in all of these models the cardiac repolarization is impaired and associated with the development of cardiac hypertrophy, heart failure and arrhythmias. Cardiac repolarization results from a concerted balance of depolarizing inward currents (mainly Ca^2+^ currents) and repolarizing outward currents (mainly K^+^ currents) in ventricular cells. Strong evidences for a circadian pattern of cardiac K^+^ channels have been provided both in rodent and human heart^19, 23, 24, 27–30^. Only a couple of controversial studies have assessed the circadian variation the ventricular L-type Ca^2+^ channel (LTCC)^31–35^, primarily composed of the Ca_V_1.2 pore-forming subunit (encoded by *CACNA1C*), Ca_V_β2 and Ca_V_α2δ1 auxiliary subunits (encoded by *CACNB2* and *CACNA2D1*, respectively)^36^. However, the interbreeding of circadian gene datasets for transcripts of *CACNA1C*, *CACNB2* and/or *CACNA2D1* in the heart show circadian expression pattern^16, 17, 19, 21, 30^. Therefore, it is reasonable to hypothesize a circadian regulation of ventricular LTCCs but the underlying mechanism remains elusive.

Herein, we investigated both *in vivo* and *in vitro* whether the cardiac LTCC oscillated daily using a unique transgenic model that expresses the luciferase reporter under the control of the cardiac *Cacna1c* promoter ^37^.

## Methods

Full methods are provided in the Supplemental Material.

## Results

### In vivo diurnal variation of cardiac Ca_V_1.2 expression

To interrogate the diurnal profile of cardiac LTCC expression, we used PCa-luc mice carrying a firefly luciferase gene under the control of cardiac *Cacna1c* rat promoter^37^. After a 150 μg/g luciferin injection, the noninvasive *in vivo* bioluminescence imaging of anesthetized PCa-luc mouse showed a strong bioluminescence signal in the heart region (Fig.1A). Although in some mice a weaker signal might be also observed in abdomen and thighs regions, this result illustrated the cardiac specificity of the *Cacna1c* promoter used. Temporal dynamics analysis in same mouse every 3 h starting at ZT0 (lights on 8 a.m.) under normal 12:12h light-dark cycle (LD cycle, Fig.1A) revealed oscillations of bioluminescence intensity (BLI) during 24 h. One might visualize a first late evening peak (inactive light-on period, ZT 9 corresponding to 5 p.m.) and a second peak at early morning (active lights-off period, ZT 21 or 5 a.m.). These experiments were repeated on 9 different 3-month old mice (Fig.1B). Quantitative analysis of the relative BLI (RLU) within the cardiac ROI showed a non-uniform distribution throughout the 24-h cycle (P_ANOVA_ < 0.05) with a 3.78 fold changes of BLI over all times. To infer whether *in vivo Cacna1c* promoter activity showed periodicity, the data were analyzed using JTK-Cycle^38^ with period set at 24 h. We found that BLI varied in a periodic manner (P_JTK_ < 0.05, phase at 19.5 and peak-to-through amplitude of 34.14%). For visualization of the temporal relationship, data were represented with harmonic cosinor model with 24 h periodicity, amplitude and phase assigned by JTK-Cycle obtained values. This graphical representation exhibits a bimodal daily pattern: a first peak at early morning (active lights-off period, ZT 19.5 or 4 a.m.) and a second smaller peak at late evening (inactive light-on period, ZT 9 corresponding to 5 p.m.).

**Figure 1.**
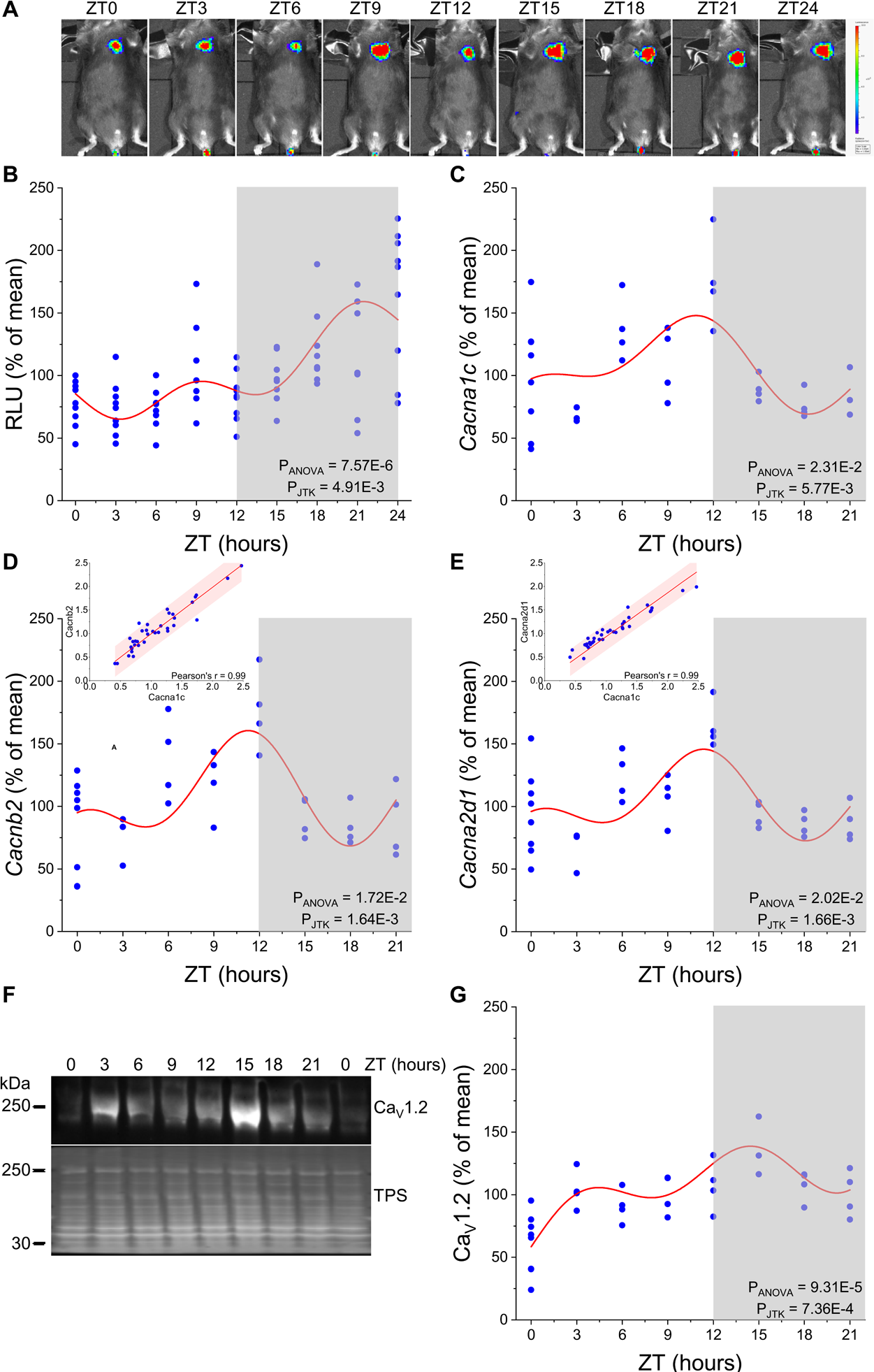
Periodic variations of Ca_V_1.2 expression in the mouse heart ventricle. **A**. Bioluminescence images of a PCa-luc mouse recorded at 3-h interval during 24 h from 0 Zeitgeber times (ZT) corresponding to light-on at 8 a.m. The color scale shows relative signal intensity or radiance (expressed in p/sec/cm^2^/sr), red being the most intense and blue the least intense. **B**. Temporal pattern of heart luciferase activities expressed as relative light units (RLU) of 9 PCa-luc mice. For each mouse at each time point, the total bioluminescence intensity counted for 16 min in the heart region of interest (ROI) were quantified, corrected from the background in the abdomen ROI and normalized to the mean. **C** to **D**. Time-dependent variations of *Cacna1c* (**C**), *Cacnb2* (**D**) and *Cacna2d1* (**E**) mRNA levels in mouse ventricular tissues. At each time point, tissues were collected on 4 mice (8 at ZT 0) and the relative mRNA levels of each transcript were normalized to the quantity RPL-32 housekeeping gene (ΔCq) and to the mean over all time (ΔΔCq). The fold of change 2^-ΔΔCq^ are expressed as %. Insets in **D** and **E**, represent the fold of change of *Cacnb2* and *Cacna2d1* transcripts, respectively, in function of fold of change *Cacna1c* mRNA in each tissue; the linear fit curves (line in red and 95% confidence level in light red) with correlation Person’s coefficients are represented. **F**. Immunoblot with Ca_V_1.2 antibody and corresponding total proteins fluorescent staining (TPS) in ventricular heart tissues from different mice collected at 3-h interval during 24 h. **G**. Time series of Ca_V_1.2 expression ratios normalized by TPS and mean of each gel in ventricular tissues from mouse heart (n=4 to 8 for each time points). Immunoblots were analyzed by using ImageJ. In **B**, **C**, **D**, **E** and **G** the results of analysis with a non-parametric one-way ANOVA (with repeated measure in B) after Aligned Rank Transform procedure with ARTool R package (P_ANOVA_) and with JTK-Cycle (P_JTK_) after being adjusted with Benjamini and Hochberg method are shown and the red lines represent the harmonic cosinor curves. For all the experiments, mice are maintained under normal 12:12h light-dark cycle, the time of light-off is shown in gray.

Taking into account the 1 to 4 h half-life of luciferase *in vivo*^39^, one might suggest that the observed promoter activity periodicity could be due to luciferase turnover. We then carried out quantitative reverse transcription polymerase chain reaction (RT-qPCR) analyses of *Cacna1c*, *Cacnb2* and *Cacna2d1* mRNAs extracted from ventricular heart tissues of mice kept in LD cycle and collected at 3-h interval during 24 h (4 mice at each time, Fig. 1C to E). We can infer that the mRNA levels fluctuated throughout the day (2.59, 1.48 and 2.19 fold changes for *Cacna1c*, *Cacnb*2 and *Cacna2d1*, respectively) showing a diurnal periodicity with a peak-to-through amplitude of 60.22% for *Cacna1c*, 61.12% for *Cacnb2* and 48.06% for *Cacna2d1* and phases at ZT12. The harmonic cosinor waveform is doubled peaked at ZT12 and between ZT21 and 0. Remarkably, we noticed a direct linear correlation between mRNA levels of the main and auxiliary subunits (insets Fig.1D and E) consistent with a coordinated control of their expressions^37^.

The presence of a transcript does not necessarily imply the presence of a functional protein. Therefore, we examined the temporal profile of the Ca_V_1.2 protein levels in the same ventricular tissue samples used for qPCR analysis. The immunoblot analysis (Fig.1F and G) documented a diurnal periodicity of Ca_V_1.2 protein levels. A 2.99-fold difference by time was observed and a periodicity characterized with a peak-to-through amplitude of 39.36% and a 1.5-h phase shift compared with *Cacna1c* mRNA. With the harmonic cosinor model, a major peak in the dark cycle (ZT15) and a weaker on in the light cycle (between ZT3 and 6) were observed.

These results suggested *in vivo* temporal differences of ventricular Ca_V_1.2 expressions throughout 24 h controlled at the transcriptional level.

### In vitro circadian modulation of Ca_V_1.2 expression and activities

Diurnal variations in ventricular Ca_V_1.2 expressions *in vivo* could be due to nervous, hormonal signals and/or circadian clock intrinsic to the cardiomyocyte. To address whether periodic oscillation of Ca_V_1.2 expression occurs by cell-autonomous mechanisms, we used isolated ventricular cardiomyocytes from 9 PCa-luc mice kept in culture during 24 h. After isolation (at ZT0) and 2 h-plating, the ventricular cardiomyocytes were acutely submitted to 50% serum shock synchronization for 2 h (given the 0 circadian time)^40^, then collected at 3-h intervals during 24 h and stored at -80°C before protein extraction. Figures 2A and B illustrate that periodic luciferase activities and Ca_V_1.2 expressions (in the same samples) persist in isolated ventricular cardiomyocytes with 24-h oscillations. Robust differences were observed over 24-h period (2.05- and 2.57-fold changes for luciferase activities and Ca_V_1.2 protein levels, respectively). JTK-Cycle analysis showed phases at 16.5 and 9 circadian time (CT) and peak-to-through amplitudes of 16.96% and 26.48%, for luciferase activities and Ca_V_1.2 protein levels, respectively. Similar to *in vivo* analysis although of lower amplitude, the *in vitro* temporal relationship of promoter activity peaked at CT6 and 19 whereas Ca_V_1.2 protein levels peaked at CT9 and 21.

**Figure 2.**
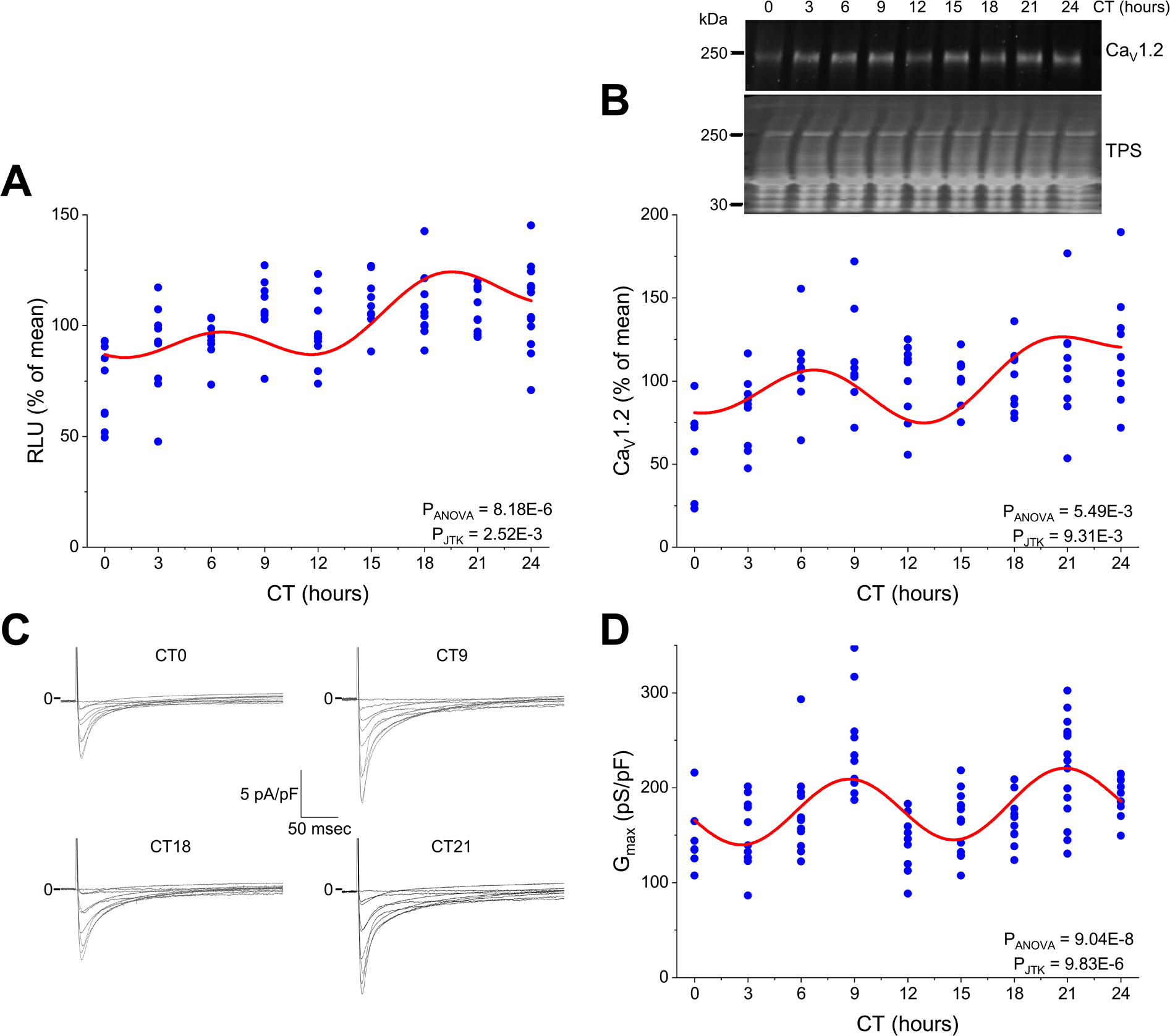
Temporal pattern of Ca_V_1.2 in isolated adult ventricular cardiomyocytes. **A**. Circadian fluctuation of luciferase activities (RLU) in cellular homogenates from isolated adult PCa-luc mouse ventricular cardiomyocytes (AMVC) maintained in culture during 24 h and sampled at 3-h interval after 50% serum shock synchronization (O Circadian time, CT0). At each time point, the measured BLI per μg of protein are expressed as the ratio of mean in each culture (n=9). **B**. Ca_V_1.2 immunoblot and TPS of AMVC from the same PCa-luc mouse collected at indicated time (upper panel) and cumulative data (lower panel) from the same samples as in A. Quantification are done as in Figure 1. **C**. I_CaL_ density traces recorded in different AMVCs maintained in culture during indicated time after serum shock synchronization (CT0). I_CaL_ amplitudes were measured and normalized to cell capacitance. **D**. 24-h temporal pattern of maximal conductance of I_CaL_ (G_max_, n=9 to 15 cells per time points). AVMCs were treated with a 2-h serum shock, then I_CaL_ were recorded every 3 h ± 15 min over a 24-h period. The adjusted p-values (from non-parametric ANOVA, P_ANOVA_ and JTK-Cycle, P_JTK_) are shown. The data were represented with harmonic cosinor shown as a solid red line.

To explore the functional consequences of these circadian rhythmicities, we analyzed the temporal pattern of LTCC currents (I_CaL_) in isolated ventricular cardiomyocytes submitted to the same protocol using whole-cell patch clamp technique. Figure 2C shows examples of the voltage- and time dependent I_CaL_ densities recorded in myocytes at indicated CTs. A large daily double peaked variations at CT9 and 21 were observed. The analysis was conducted on 7 to 15 different myocytes from 8 mice at 3-h interval (± 15 min) during 24 h and current-voltage relationships were fitted with a function combining the Goldman-Hodgkin-Katz equation and the Boltzmann relationship to estimate maximal conductance (G_max_)^41^. The data presented in Figure 2B revealed a 1.71-fold oscillation of G_max_ over 24 h. JTK-cycle analysis identified G_max_ periodic variations (phase at CT9 and peak-to-through amplitude of 38.7%) in concordance with Ca_V_1.2 protein expression (Fig.2B), with peaks at CT9 and 21. Moreover and consistent with coordinated expression of LTCC subunits *in vivo* (Fig.1C and D), neither the time-nor the voltage-dependent properties of I_CaL_ showed variations during 24 h (Table 1).

**Table 1.** Temporal cellular electrophysiology characteristics of isolated ventricular cardiomyocytes.

|  | n<sub>cell</sub> | CT0<br>8 | CT3<br>10 | CT6<br>13 | CT9<br>10 | CT12<br>9 | CT15<br>13 | CT18<br>10 | CT21<br>15 | CT24<br>12 | P<sub>ANOVA</sub> |
| --- | --- | --- | --- | --- | --- | --- | --- | --- | --- | --- | --- |
| C<sub>m</sub> (pF) |  | 156.67<br>[135.9,177.4] | 147.49<br>[134.7,169.7] | 145.43<br>[132.4,160.2] | 140.90<br>[97.6,163.3] | 150.65<br>[98.1,220.7] | 150.64<br>[135.1,181.4] | 126.07<br>[120.2,171.1] | 102.88<br>[95.2,144.2] | 106.19<br>[101.7,156.7] | 0.155 |
| Activation |  |  |  |  |  |  |  |  |  |  |  |
| T<sub>peak</sub> (msec) |  | 8.75<br>[7.7,12.2] | 8.80<br>[8.3,10.1] | 9.20<br>[8.8, 9.4] | 7.80<br>[7.5, 9.0] | 9.00<br>[8.4, 11.9] | 8.20<br>[7.4, 8.4] | 8.65<br>[7.4, 9.8] | 7.70<br>[7.2, 8.8] | 7.40<br>[6.5, 8.2] | 0.245 |
| V<sub>50</sub> (mV) |  | -9.08<br>[-11.2, -8.4] | -11.98<br>[-14.8, -8.3] | -11.87<br>[-12.4, -9.3] | -10.26<br>[-11.7, -8.9] | -12.79<br>[-13.5, 9.5] | -10.65<br>[-14.1, -8.5] | -10.40<br>[-1.8, -7.8] | -11.65<br>[-13.3, -10.2] | -10.39<br>[-12.0, -8.9] | 0.415 |
| k (mV) |  | -7.35<br>[-8.2, -6.6] | -5.93<br>[-7.9, -5.1] | -5.61<br>[-6.4, -5.4] | -6.21<br>[-7.7, -5.8] | -5.27<br>[-6.6, -4.8] | -5.97<br>[-6.6, -5.3] | -6.09<br>[-6.9, -5.8] | -5.78<br>[-6.8, -5.2] | -6.71<br>[-7.8, -5.4] | 0.150 |
| Inactivation |  |  |  |  |  |  |  |  |  |  |  |
| V<sub>50</sub> (mV) |  | -23.94<br>[-30.3, -22.1] | -26.08<br>[-26.9, -22.4] | -24.05<br>[-25.3, -23.7] | -23.77<br>[-25.3, -23.5] | -26.27<br>[-27.2, -25.2] | -25.70<br>[-29.2, -24.2] | -24.39<br>[-25.6, -22.1] | -25.83<br>[-27.5, -24.4] | -25.89<br>[-30.5, -22.8] | 0.561 |
| k (mV) |  | 6.79<br>[5.4, 8.5] | 5.75<br>[4.9, 5.9] | 5.78<br>[5.5, 6.4] | 6.86<br>[6.2, 7.2] | 5.50<br>[5.4, 6.1] | 5.87<br>[5.5, 6.3] | 6.43<br>[5.6, 7.8] | 6.19<br>[5.7, 6.9] | 7.01<br>[6.2, 8.3] | 0.154 |
| τ<sub>slow</sub> (msec) |  | 90.14<br>[83.5, 104.5] | 78.52<br>[70.8, 85.7] | 88.36<br>[80.4, 94.1] | 89.62<br>[78.5, 101.5] | 86.59<br>[84.2, 90.3] | 84.22<br>[76.6, 96.3] | 87.32<br>[79.2, 121.9] | 81.73<br>[75.6, 95.7] | 90.96<br>[84.7, 128.6] | 0.433 |
| τ<sub>fast</sub> (msec) |  | 9.20<br>[7.1, 10.1] | 9.57<br>[8.9, 11.7] | 9.79<br>[8.4, 11.2] | 8.19<br>[7.1, 9.1] | 9.26<br>[6.6, 10.3] | 10.56<br>[8.1, 11.3] | 8.14<br>[7.3, 9.8] | 8.94<br>[8.1, 10.2] | 7.05<br>[6.8, 9.4] | 0.209 |

Data are presented as median [95% confidence interval]. n<sub>cell</sub> is the number of cell analyzed. T<sub>peak</sub> is the time from the onset of the voltage step to the peak of current at 0 mV, V<sub>50</sub> is the potential of half-maximum and k is the slope factor for both activation and inactivation. τ are the time constants of the two components of the time course of inactivation at 0 mV subscripted fast and slow, respectively. P<sub>ANOVA</sub> are the p values of one-way ANOVA on aligned rank transform data with the ARTool R package.

The *in vitro* persistent periodicity of transcriptional control of ventricular Ca_V_1.2 activity and expression supports the involvement of endogenous cellular clock mechanism.

### Absence of diurnal variation of Cav1.2 in atrial heart tissues

During the *in vivo* analysis of diurnal variation of *Cacna1c* in ventricular tissues, we also collected in the same mice the atrial tissues. Surprisingly, the analysis of mRNA levels of *Cacna1c*, *Cacnb2* and *Cacna2d1* did not show diurnal periodicity in atria (Fig.3).

**Figure 3.**
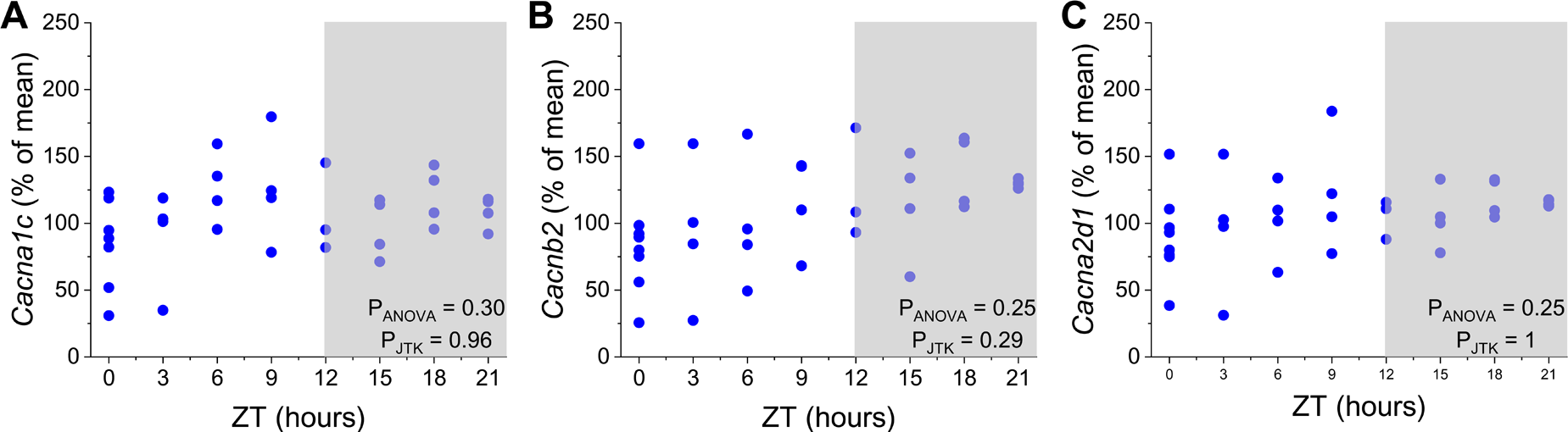
No change of atrial LTCC transcripts as a function of the time of the day. Expression of *Cacna1c* (**A**), *Cacnb2* (**B**) and *Cacna2d1* (**C**) transcripts in atrial heart tissues of mice maintained under normal 12:12h light-dark cycle and collected at 3-h interval during 24-h cycle (n=4 to 8 for each time points). The adjusted p-values (from non-parametric ANOVA, P_ANOVA_ and JTK-Cycle, P_JTK_) are shown.

To analyze the presence of a molecular clock in our samples, we measured by qPCR sequential mRNA levels of major clock components including transcription factors *Bmal1* (Fig.4A), *Clock* (Fig.4B), *Per1* (Fig.4C), *Per2* (Fig.4D), *Cry1* (Fig.4E), *Cry2* (Fig.4F), *Dbp* (Fig.4G), *Nfil3* (Fig.4H), *Nr1d1* (Fig.4I) and *Nr1f1* (Fig.4J) in both ventricular (left panels) and atrial (right panels) tissues collected during 24 h. As previously reported^16–19, 22, 23, 42^, all components of the circadian clock transcripts were expressed in ventricular and atrial mouse tissues and showed day-night rhythms except *Nr1f1* (encoding for RORα). In ventricular tissues (Fig.4J left panel), *Nr1f1* mRNA levels showed 1.79-fold variations over all time and periodically fluctuated (phase at CT9 and peak-to-through amplitude of 40.87%) with a temporal cosinor pattern peaking between CT6-9 and CT15-18. In contrast, no significant difference in the level of *Nr1f1* transcript expression were observed during 24-h cycle in atrial tissues (Fig.4J right panel). There were no differences in the level of RORα protein expression between ventricular and atrial heart tissues, when compared at the same time points (supplemental Fig.1).

**Figure 4.**
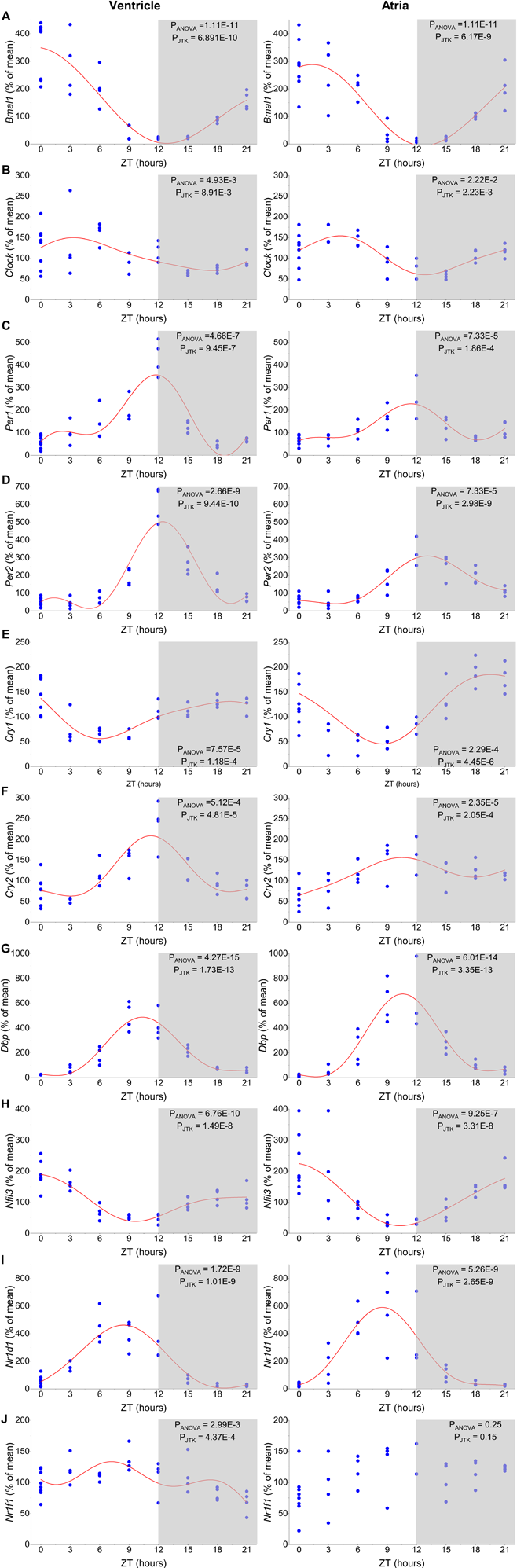
Diurnal expression of core clock genes in ventricular and atrial heart tissues at 8 different time points. Expression of clock transcripts across the 24 h cycle in ventricular (left panels) and atrial (right panels) heart tissues from the same mice (n=4 to 8 for each time points and group) maintained under normal 12:12h light-dark cycle. The adjusted p-value (from non-parametric ANOVA, P_ANOVA_ and JTK-Cycle, P_JTK_) are shown. The data are represented with harmonic cosinor when p-values <0.05 and shown as a solid red line.

These results point out that there is a functional circadian clock in mouse ventricular and atrial tissues and suggested that RORα could be an important factor responsible for the diurnal rhythm of ventricular LTCC expressions.

### RORα regulates circadian expression of ventricular Ca_V_1.2

To explore mechanism through which the circadian clock regulated periodic expression of Ca_V_1.2 in the ventricles, we first analyzed the temporal patterns of the RORα protein levels. Consistent with qPCR analysis (Fig.4J), we denoted that RORα protein levels both *in vivo* (2.69-fold change, Fig. 5A) and *in vitro* (4.63-fold change, Fig.5B) varied periodically (phases at ZT13.5 and CT12, peak-to-through amplitude of 36.08 and 72.42%, for *in vivo* and *in vitro* respectively) over the course of the 24-h cycle peaking at 7.5 and 18h.

**Figure 5.**
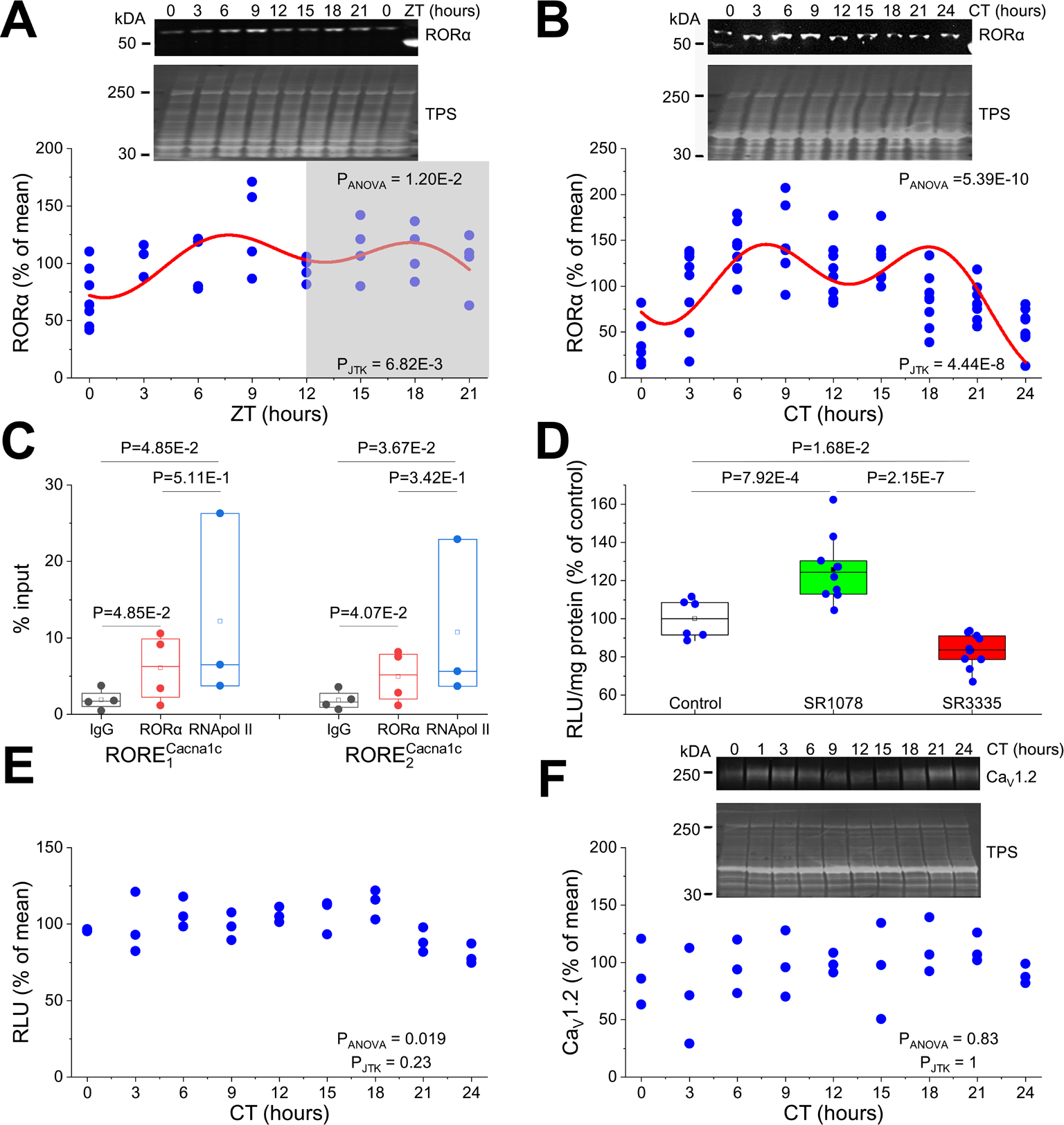
RORα mediates circadian changes in ventricular Cav1.2. **A** and **B**. Immunoblot with RORα antibody and TPS (top panels) and quantifications following band densitometry (lower panels) assessing RORα protein levels in ventricular heart tissues (**A**, n=4 to 8 for each time points) and isolated AVMCs (**B**, n=8 for each time points) collected at 3-h interval during 24 h. **C**. ChIP assay of immunoprecipitated chromatin from AVMCs (4 different isolations) with a RORα antibody and quantified qPCR, using a primer sets specific for the putative ROREs in the cardiac *Cacna1c* mouse promoter. Non-immune IgG and DNA immunoprecipitated with RNA polymerase II antibody served as negative and positive controls respectively. **D**. Luciferase activities of isolated AVMCs from PCa-luc mice treated for 24 h in the absence (control, white box, n=10) or in the presence of either RORα agonist, 10-μM SR1078 (green box, n=10), or RORα inverse agonist, 5-μM SR3335 (red box, n=10). Data are presented as median (line) and 95% confidence interval (box). **E** and **F**. Time-course data of luciferase activities (**E**) and Ca_V_1.2 protein levels (**F**, upper inset showing immunoblot with Ca_V_1.2 antibody and TPS) in AVMCs maintained during 24 h in presence of 5-μM SR3335 (n=3 for each time points). The adjusted p-value (from non-parametric ANOVA, P_ANOVA_ and JTK-Cycle, P_JTK_) are shown and overlay of harmonic cosinor models are shown when p-values <0.05.

*In silico* examination of the ∼4 kb cardiac promoter region of mouse *Cacna1c* using TFsearch (https://alggen.lsi.upc.es/cgi-bin/promo_v3/promo/promoinit.cgi?dirDB=TF_8.3), revealed 2 canonical consensus binding sites for RORα: RORE^Cacnalc^ TGGCAGGTCA^-848^ and RORE^Cacnalc^ TGTAAGGTCA^-1030^ (site locations relative to the transcription start site of Mus musculus strain C57BL/6J NC_000072.6:c119200736-119196242). To examine the potential roles of these ROREs in RORα mediated regulation of *Cacna1c* expression, we conducted ChIP followed by qPCR on mouse ventricular cardiomyocyte freshly isolated at ZT 9 using (n=4 independent cell isolations) using either mouse anti-RORα4 antibody^26^, anti-RNA Polymerase II Antibody (RNA PolII) as positive control or normal Rabbit IgG as negative IP control (Figure 5C). Relative to the IgG, we found ∼3-fold enrichment of RORα on the two ROREs.

We next examined the effect of pharmacological activation and inhibition of RORα *in vitro*. After serum synchronization, isolated ventricular myocytes from 3 PCa-luc mice were kept in culture in the absence or the presence of either SR1078, a specific RORα agonist^43^ or SR3335, a RORα inverse agonist^44^ and luciferase activities were measured 24 h later. As shown in Fig. 5D, SR1078-(10 μM, green box) and SR3335-treament (5 μM, red box) increased and reduced, respectively, *Cacna1c* promoter activity. In addition, we showed that in presence of 5-μM SR3335, *Cacna1c* promoter activities (Fig.5E) and Ca_V_1.2 protein levels (Fig.5F) were devoid of any discernable periodicity over the course of the 24-h cycle.

These data indicate that the ventricular Ca_V_1.2 24-h periodic expression is controlled at transcriptional level by RORα.

### RORα as a major component in control of the diurnal cardiac repolarization periodicity

It has been reported in rodents that the major early repolarizing K^+^ current formed by Kv4.2/4.3 and KChIP2 subunits showed circadian variation of expression^23, 27^. Even if it has been shown that the Kidney-Enriched Krueppel-Like Factor 15 (Klf15) transcriptionally controls periodic expression of KChIP2, the molecular mechanism controlling Kv4.2/4.3 circadian rhythmicity is still elusive^23^. We analyzed by qPCR the diurnal pattern of *Klf15*, *Kcnip2*, *Kcnd2* and *Kcnd3* (encoding Kchip2, Kv4.2 and 4.3, respectively) transcripts in ventricular (Fig.6 left panels) and atrial (Fig.6 right panels) heart tissues of the same mice. Consistent with previous report^23^, we observed a periodic expression of *Kcnip2* and *Klf15* mRNA in both cardiac tissues (Fig.6A and B). However, as shown in Figure 6C and D, the diurnal variations of *Kcnd2* and *3* (2.16- and 2.01-fold change, respectively) observed in ventricles (phase at CT6, peak-to-through amplitude of 64.20 and 53.49%, for Kcnd2 and Kcnd3, respectively) were absent in atrial tissues of the same mice, suggestive of a potential involvement of RORα. Indeed, in isolated ventricular cardiomyocytes the 2.12-fold variations of Kv4.3 protein levels throughout the 24-h cycle (phase at CT9, peak-to-through amplitude of 44.588%, Fig. 6E) was blunted in the presence of SR3335 (Fig. 6F). The analysis of mouse *Kcnd2* and *Kcnd3* promoters pointed out the presence of several ROREs: RORE*^Kcnd2^*: AGAAAGGTCA^-923^ [NM_019697]; 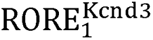 TGACCTTCTG^-778^ and 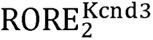 TGACCTCAAA^-3008^ [NM_019931]). ChIP assay (Fig.6 G) showed ∼3-fold enrichment for RORα binding in the RORE^Kcnd2^ and 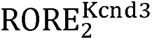 but no enrichment in the 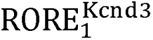 regions.

**Figure 6.**
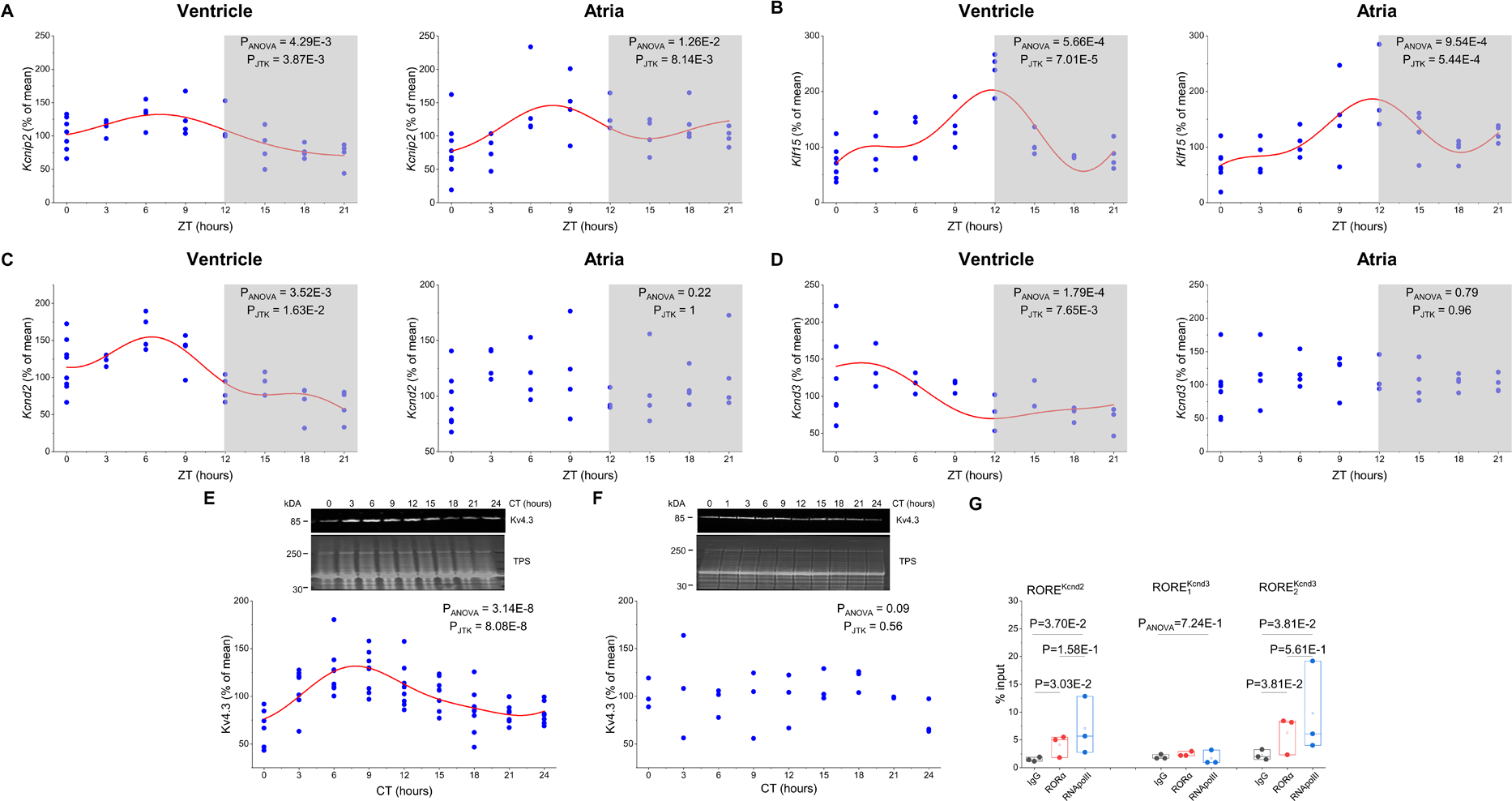
Potential role of RORα in mediating diurnal variations in transient K^+^ outward channel transcripts. **A** to **D**. Expression of *Kcnip2* (**A**), *Klf15* (**B**), *Kcnd2* (**C**) and *Kcnd3* (**D**) transcripts across the 24 h cycle in ventricular (left panels) and atrial (right panels) heart tissues from the same mice (same sample as Fig.1 and 3) maintained under normal 12:12h light-dark cycle. **E** and **F**. Time-course of Kv4.3 protein levels in AVMCs maintained during 24 h in the absence (**E**, n= 8 for each time points) or in the presence of 5-μM SR3335 (**F**, n= 3 for each time points). Upper insets showing immunoblot with Kv4.3 antibody and TPS. The adjusted p-value (from non-parametric ANOVA, P_ANOVA_ and JTK-Cycle, P_JTK_) are shown and overlay of harmonic cosinor models are shown when p-values <0.05 and shown as a solid red line. **G**. ChIP assay of immunoprecipitated chromatin from AVMCs (4 different isolations) with a RORα antibody and quantified qPCR, using a primer sets specific for the putative ROREs in the *Kcnd2* and *3* mouse promoters. Non-immune IgG and DNA immunoprecipitated with RNA polymerase II antibody served as negative and positive controls respectively.

## Discussion

Our study shows that the LTCC expression presents diurnal variations and demonstrated the pivotal role of the transcription factor RORα, as proximal mediator of cardiac biological clock in this circadian regulation, which might have potential pathophysiological consequences.

Even if diurnal pattern of LTCCs have been previously proposed in the heart, the results are somewhat inconsistent. In ventricular cardiomyocytes from Wistar rats, a nocturnal specie, the I_CaL_ measured at two times of the day was higher in active period (ZT15), even though the Ca_V_1.2 mRNA and protein levels did not change^31, 34^. In ventricular myocytes of diurnal guinea-pig, a *Clock*-*Bmal1*-dependant circadian variation of Ca_V_1.2 proteins through PI3K-Akt pathway was associated with simultaneous larger I_CaL_ at ZT3 than at ZT15, but without diurnal variations of *Cacna1c* transcripts^35^. Conversely, Ca_V_1.2 currents, mRNA and protein expressions peaked during the night under the control of Erk- and PI3K-Akt signaling pathways in embryo chick hearts^32^ but mRNA and protein expressions of Ca_V_1.2 in WT and cardiomyocyte-specific *Clock* mutant mouse heart remained constant throughout the day^33^ whereas these pathways have been shown to regulate cardiac LTCCs^43, 45^. Our results showed both *in vivo* and *in vitro* in a mouse model (Fig. 1 and 2) consistent diurnal variations of cardiac *Cacna1c* promoter during 24-h cycle, leading to similar periodic variations of Ca_V_1.2 transcripts (with a 3-h phase shift) that precede (by 1.5 h) the simultaneous circadian oscillations of Ca_V_1.2 proteins and currents. While several factors could generate such disparities (species, age, sampling interval, *in vivo*/*in vitro*, culture conditions….), our data echoed the results of unbiased approaches by microarrays and RNAseq showing circadian expression of LTCC transcripts. Notably, mouse circadian transcriptomic datasets assessed a biphasic circadian pattern of heart *Cacna1c*, *Cacnb2* and/or *Cacna2d1* genes, with one expression peak in the light phase, and a second peak in the dark^16, 17, 19^, in agreement with our results. Likewise, query of the primate library of diurnal transcriptome revealed that *CACNB2* and *CACNA2D1* genes in heart peaked in the morning^21^. Moreover, a recent analysis by qPCR in failing human ventricular heart tissues^30^ reported a circadian pattern of *CACNA1C* expression with 54.1% variance ascribed to a 24 h circadian pattern. In addition, a search in database of cardiomyocyte-specific circadian gene knock-out model revealed alterations of LTCC transcript levels^22, 46, 47^, pointing out a circadian clock gene-dependent regulation.

In contrast to ventricular tissues, the LTCC and *Nr1f1* transcripts failed to oscillate in mouse atrial tissues (Fig. 3 and 4), whereas cyclic variations of other major components of the circadian clock were maintained. Consistent with these results, a human tissue sample database lookup^48^ revealed that *CACNA1C* and *CACNA2D1* gene in heart atrial do not met criteria for periodicity. This leads us to hypothesize that RORα was involved in ventricular Ca_V_1.2 periodicity. Indeed, we observed that specific RORα inverse agonist prevented circadian rhythmicity of both promoter activities and Ca_V_1.2 protein levels. Analyzing the cardiac promoter region of *Cacna1c* we found 2 putative ROREs. ChIP assay showed that RORα is indeed recruited onto the cardiac *Cacnac1c* promoter in mouse ventricular cells. Matching ROREs in the same sequential order were also found in the upstream region of *Cacna1c* orthologues from rat and human (TGGCAGGTCA^-840^ and TGGAAGGTCA^-926^ for Rattus norvegicus NC_005103.4:c151273977-151269116 and CTTGAGGTCA^-2640^ and CGTGAGGTCA^-3892^ for Homo sapiens NC_000012.12:1965780-1970880) providing strong evidence for similar and specific gene regulation. Similarly, in mouse brain tissue that expressed the short N-terminal Ca_V_1.2 isoform (Ca_V_1.2-SNT) driven by a different promoter^37^, 4 putative ROREs have been characterized^49^. This study pointed out a repressive Rev-Erbα-dependent circadian rhythmic Ca_V_1.2-SNT expression participating in setting the SCN circadian clock during the late night, but also showed a RORα-induced Ca_V_1.2 promoter activity. Indeed, RORα competes with Rev-Erbα for binding of their shared DNA binding elements and their opposing activities are important in the maintenance of the core mammalian circadian clock^50^. Schmutz et al^49^ observed that circadian amplitudes of Ca_V_1.2-SNT expression were only attenuated by the loss of Rev-Erbα, suggestive of potential role of RORα. We thus propose RORα as output mediator of the circadian molecular clock in the Cav1.2 diurnal regulation.

Recently, the function of RORα in the heart has gained peculiar interest due to its potential cardioprotective role in ischemia/reperfusion, hypertrophy development and diabetic cardiomyopathy, which could also relay melatonin beneficial cardiac effects^26, 51–53^. However, to the best of our knowledge, RORα role in cardiac circadian clock has not been investigated. Nevertheless, it has been shown that thyroid status, which impact cardiac repolarization through regulation of K^+^ and Ca_V_1.2 channels^54^, altered the 24-h rhythms in circadian clock, notably of RORα, in rat heart^55^. As for Ca_V_1.2, we observed that the diurnal oscillations of the pore forming subunits of K^+^ channels (Kv4.2 and 4.3), underlying the cardiac transient K^+^ outward current (I_to_), were lost in atrial heart tissues (Fig. 6). Analysis of Mus musculus *Kcnd2* promoter found the presence of one RORE in the mouse, which is conserved in rat and human promoters (at position-934 and -391 for Rattus norvegicus [NC_051339.1] and for Homo sapiens [NM_012281], respectively). For *Kcnd3,* only matching RORE^Kcnd3^ are present in rat and human promoter (TGACCTCAGA^-3288^ for Rattus norvegicus [NM_0311739] and TGACCTTCCC^-1295^ for Homo sapiens [NM_172198]). Consistently, RORα bound to RORE^Kcnd2^ and 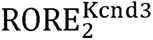 but not on 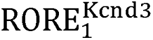. Taken together, this suggests a conservative RORα-dependent mechanism for the circadian control of cardiac repolarization.

Although, changes in I_to_ do not produce much change in ventricular repolarization in humans, it is quite possible that the additive diurnal changes in Ca_V_1.2 expression, could contribute to the production of potentially lethal arrhythmias. Indeed, Ca_V_1.2 current increase and concomitant K^+^ current decrease lead to prolongation of cardiac repolarization, which allows recovery of Ca_V_1.2 inactivation and generation of an inward depolarizing current during repolarization that triggers early after depolarization that might initiate sustained arrhythmias by reentrant mechanisms^36^. The temporal occurrence of ventricular arrhythmias is known to have the first peak between morning and noon and the second peak in the evening^3–5^, which could be related to the increases on Ca_V_1.2 at both subjective light- and dark-times observed herein.

In conclusion, our results uncovered a new circadian RORα-dependent cardiac repolarization regulatory mechanism that, in addition to neuronal and hormonal influence, could participate to the daily periodic control of cardiac events.

## Nonstandard Abbreviations and Acronyms

AMVC: adult mouse ventricular cardiomyocytes
BLI: bioluminescence intensity
BMAL1: Brain and muscle ARNT-like protein-1
CACNA1C: Ca2^+^ voltage-gated channel subunit alpha1 C gene
CACNB2: Ca2^+^ voltage-gated channel auxiliary subunit beta 2 gene
CACNA2D1: Ca2^+^ voltage-gated channel auxiliary subunit alpha2delta 1
Cav1.2: Ca2^+^ channel, voltage-dependent, L type, alpha 1C
CaV1.2-SNT: short N-terminal CaV1.2 isoform
ChIP: Chromatin immunoprecipitation assay
CLOCK: Circadian locomotor output cycles kaput
CRY: Cryptochrome
CT: circadian time
DBP: D-site albumin-binding protein
G_max_: maximal conductance
I_CaL_: LTCC currents
Kv4.2/Kv4.3: Pore forming proteins of the K^+^ transient outward channel
KCND2: K^+^ voltage-gated channel subfamily D member 2 gene
KCND3: K^+^ voltage-gated channel subfamily D member 2 gene
KChIP2: Kv channel-interacting protein 2
KCNIP2: K^+^ voltage-gated channel interacting protein 2 gene:
Klf15: Kidney-Enriched Krueppel-Like Factor 15
LD cycle: light-dark cycle
LTCC: L-type Ca2^+^ channel
NFIL3: Nuclear factor interleukin 3 regulated
NR1D1/2: Nuclear receptors REV-ERBα/β gene
NR1F1: Retinoid-related orphan receptor alpha gene
PCa-luc: Transgenic mouse model that expresses the luciferase reporter under the control of the cardiac *Cacna1c* promoter
PER: Period
qPCR: Quantitative polymerase chain reaction
RLU: relative BLI
ROI: Region of interest
RORα: the retinoid-related orphan receptor α
RORE: ROR response element
SCN: Hypothalamic suprachiasmatic nucleus
SR1078: RORα agonist
SR3335: RORα inverse agonist
TF: transcriptional factors
ZT: Zeitgeber time

## Acknowledgments

The authors thank Catherine Cailleau (Université Paris-Saclay, CNRS, Institut Galien Paris-Saclay) for her assistance with *in vivo* bioluminescence experiments, Patrick Lechene (Université Paris-Saclay, Inserm, Signalling and Cardiovascular Pathophysiology, UMR-S 1180) for his help with R software and the platforms from “Ingénierie et Plateformes au Service de l’Innovation Thérapeutique” University of Paris-Saclay IPSIT: the AnimEX plateform (Julie Burlot, Valérie Domergue) for animal care; the ACTAGen plateform (Claudine Deloménie) for qPCR; the CIBLOT plateform (Delphine Courilleau) for *in vitro* bioluminescence experiments.

## Author Contributions

All authors contributed to conception design and interpretation of the data. EP, GG, CP, RP and JPB contributed to acquisition and analysis of the data. AMJ provided antibody. AMG and JPB acquired the financial supports. All authors read, corrected and approved the final article.

