## Supplemental material for "Circadian regulation of Ca_V_1.2 expression by RORα in the mouse heart"

**Methods**

All experiments were carried out according to the ethical principles laid down by the French Ministry of Agriculture (agreement B9201901) and were performed conform to the guidelines from Directive 2010/63/EU of the European Parliament on the protection of animals after positive feedback of the local Ethics Committee (APAFIS#1296-2015072211438825v3).

*Mouse model and cardiomyocyte culture*

Experiments were performed with 3-month aged C57BL/6 N (Charles River) and PCa-luc male mice^37^ maintained under normal 12:12h light-dark cycle (light on 8 a.m.) with free access to food and water. Hearts were collected from Na^+^-pentobarbital (100 mg/kg, i.p. complemented with 1000U/kg heparin) anesthetized mice, washed in ice-cold (0 °C) oxygenated Tyrode’s solution (in mmol/L: NaCl 113, KCl 4.7, MgSO_4_ 1.2, KH_2_PO_4_ 0.6, NaH_2_PO_4_ 0.6, NaHCO_3_ 1.6, HEPES 10, Taurine 30, D-glucose 20, Insulin 0.1 µg/ml adjusted to pH 7.4) and either directly frozen nitrogen (ventricular and atrial tissue separately) and kept at -80°C or used for ventricular cardiomyocyte isolation as previously described^37^. 25 000 to 50 000 isolated mouse ventricular cardiomyocytes per well were seeding for 2 h on plastic dishes coated with laminin (Gibco™, 23017015), synchronized by 50% Fetal Bovine Serum shock (Gibco™, 10500064) for 2 h, and then maintained in culture for up to 24 h at 37°C-5 % CO_2_ atmosphere in Tyrode’s solution supplemented with 1 mmol/L CaCl_2_, B-27™ (Gibco™, 17504044), 10 000 u/ml penicillin and 10 000 μg/ml streptomycin.

*Bioluminescence*

*In vivo* bioluminescent imaging was performed using the IVIS® Lumina III (PerkinElmer). Mice were anesthetized using 3% isoflurane (1-2 L/min O_2_) before IP injection with 150 μg/g VivoGlo™ Luciferin, In Vivo Grade (Promega, P1043) diluted in sterile PBS. Anesthesia was maintained (0.5% isoflurane + 0.5 L/min O_2_) over the course of whole ventral image acquisition at 37°C performed sequentially from 1 to 20 min post substrate injection (exposure time 110 sec every 2 min). All images were analyzed with Living image software (PerkinElmer). Region of interest (ROI) was drawn around the heart region, and average radiant efficiency was quantified (photons/sec/cm^2^/steradian). To evaluate residual background, the ROI was moved away from the mouse heart region where no signal was observed, and was used to subtract signals for calculating the relative bioluminescence intensity, which was further normalized to the mean over all time for each animal.

For *in vitro* luciferase assays, cultivated ventricular cardiomyocytes collected at each time points were lysed with 150 µl Potassium phosphate buffer Triton X-100. Protein content was determined by BCA protein assay and luminescence measured using EnVision Xcite (Perkin Elmer) in presence of 100 µl luciferin (Promega, E1960). The relative light units per µg of total proteins in the lysate were normalized to the mean over all time for each cell culture, to take into account expression variation of the transgene between mice.

*Quantitative Real-Time PCR*

The RNA extraction and RT-qPCR quantification were performed as previously described^37^. The primers are listed in Table S1. Because *Ywhaz* (14-3-3 protein zeta/ δ) and *Tbp* (TATA-box Binding Protein) showed significant variations over all 24-h cycle, only *Rpl32* (Ribosomal Protein L32) was used as housekeeping gene (HE). For each gene of interest, the difference between Cq experimental values for the gene of interest and HE amplified in the same tube (ΔCq) was first calculated, then the difference between ΔCq and the mean ΔCq for a given gene in all samples was calculated. Finally, the value of 2^−ΔΔCq^ was calculated to get the expression fold change.

*Immunoblotting*

Total protein extracts from mouse ventricular tissue samples or from cultured isolated cardiomyocyte lysates (30 μg) were resolved by SDS‐PAGE as previously described^37^ using antibodies to Ca_V_1.2 (Sigma-Aldrich, AB5156; 1:500 diluted in 5% non-fat milk) RORα (GeneTex, GTX79267; 1:1,000 diluted in 2% gelatin) and Kv4.3 (Sigma-Aldrich, P0358; 1:1,000 diluted in 2% gelatin). For loading normalization, total proteins (TPS) were stained with Revert 700 (Li‐Cor, 926-11021) and imaged with iBright FL1000 (ThermoScientific). After incubation with a peroxidase-conjugate goat anti-rabbit IgG second antibody (Santa Cruz, sc-2004; 1: 10,000), blots were revealed using luminol (SuperSignal West Pico Plus et Femto; ThermofisherTM) and recorded with the multi‐exposure mode of the iBright FL1000. The results were analyzed with Image J software and the relatives band intensities to TPS were normalized to the mean values of the same gel.

*Patch-clamp*

L-type Ca^2+^ currents (I_CaL_) were recorded at 22–24 °C using the whole-cell patch-clamp method (Axopatch-1D amplifier, Axon Instruments) with 1–1.8 MΩ micropipettes. The capacitive current was determined as previously described^41^ and the series resistance was compensated electronically (40–70 %). Electrophysiological recordings were filtered at 2 kHz, digitized and sampled at 10 kHz using a Digidata 1200 series interface and pCLAMP8 software (Axon Instruments). During experiments, cells were perfused with an external solution containing (in mmol/L) 140 NaCl, 1.1 MgCl_2_, 4 CsCl, 1.8 CaCl_2_, 10 glucose, 10 Hepes (pH 7.4), while the patch pipette was filled with a solution containing (in mmol/L): 135 CsCl, 4 MgCl_2_, 5 EGTA, 3 Na_2_-phosphocreatine, 5 Na_2_-ATP, 10 HEPES and 10 D-glucose; pH 7.2. I_CaL_ were elicited at 0.1-Hz frequency from a holding potential of -80 mV by a conventional 2 pulse protocol: a 300 msec voltage steps in 10 mV increments from -60 to +60 mV for current-voltage relationships, followed by a post voltage steps from -40 to 0 mV during 300 msec to determine the degree of inactivation. Prior to it, the voltage-gated Na^+^ channels were inactivated by a 500-msec ramp from -80 to -40 mV. The currents were normalized to the membrane capacitance and the I-V relationships were fitted with a function combining the Goldman-Hodgkin-Katz equation and the Boltzmann relationship to estimate maximal conductance: I=(V−V_rev_) G_max_ ((1+exp((V−V_0.5_)/k)) +1), where V is the voltage and the parameters estimated by the fit, where V_rev_ is the reversal potential; G_max_, the maximal conductance; V_0.5_, the half point of the relationship; and k the slope factor. The activation kinetic was measured as the time from the onset of the voltage step to the peak of current. The time course of inactivation was determined by analysis of the decay phase of current traces in response to voltage steps. Best fits were obtained with an equation including a sum of two exponentials plus a constant expressed as A_fast_ exp(-t/τ_fast_) +A_slow_ exp(-t/τ_slow_) +A_0_, where τ and A are the time constants and the amplitudes of the two components subscripted fast and slow, respectively, and A_0_ is the amplitude of the time-independent component. The current availability in function of the voltage was analyzed by fits of the amplitude of current normalized to its maximum to a Boltzmann equation.

*Chromatin immunoprecipitation assay*

ChIP assay was performed using Pierce^TM^ agarose ChIP kit (Thermo Fisher Scientific, 26156) as per manufacturer’s instruction. Mouse adult ventricular cardiomyocytes isolated at ~ZT 9 were cross-linked during 8 min in 1% formaldehyde at 25°C. After quenching with 0.125 M glycine, lysis and MNase digestion, the 75 to 200 kB DNA fragment were incubated overnight at 4°C with either mouse anti-RORα4 antibody (10 µg)^26^, anti-RNA Polymerase II Antibody (10 µL) as positive control or normal Rabbit IgG (1 µL) as negative IP control. After an incubation with ChIP Grade Protein A/G Plus Agarose for 2 h, DNA-protein complexes were eluted and the cross-linking reversed. The amount of immunoprecipitated-DNA relative to each input DNA was determined by qPCR in duplicate with primers listed in Table S1. The results were reported as percent of input 2^[ΔCq−Log^_2_^(DF)]^_*_100, where ΔCq=$C_{q}^{\mathrm{Input}}$-$C_{q}^{\mathrm{IP}}$ and DF is the dilution factor.

*Data analysis.*

Analysis were conducted using OriginPro 2022, Clampfit10.7 and R statistical computing software.

For time series analysis, we used a non-parametric factorial ANOVA after Aligned Rank Transform procedure with ARTool R package (https://github.com/mjskay/ARTool) to decipher noise from oscillation under the assumption of the null hypothesis (uniformity of the data). If this analysis ascertains lack of uniformity (P_ANOVA_<0.05, when the null hypothesis is rejected), we then pursuit to document possible rhythmicity using Jonckheere-Terpstra test, which detects monotonous trends assuming symmetric wave form^38^. When Benjamini-Hochberg corrected p-values (P_JTK_) are <0.05 we consider data as rhythmic. As negative control, we check that for the diurnal data with P_ANOVA_>0.05, the rhythmicity null hypothesis is correct (P_JTK_>0.05). We graphically represent the observed rhythmicity by fitting the data with a harmonic cosinor linear model y_0_+𝐴_1_𝑐𝑜𝑠 [(2π t/24) + φ_1_] + 𝐴_2_𝑐𝑜𝑠 [(4π t/24) + φ_2_] + 𝐶, where y_0_ is the offset, t is the time, C is an error term (assumed to be independent and normally distributed with mean zero and fixed unknown variance), A_1_, A_2_, φ_1_ and φ_2_ are the amplitudes and the acrophase of the sinusoid set at JTK values.

**Table S1.** List of primer sequences

**Supplemental Figure 1. RORα protein expressions in ventricular and atrial heart tissues**. RORα immunoblot and TPS of ventricular (V) and atrial (A) heart tissues from two mice collected at ZT2.

**Supplemental Figure 2. Uncropped gel images.**

**Table S1.** List of primer sequences

| Target gene | Sequence |
| --- | --- |
| *Cacna1c*  *Cacnb2*  *Cacna2d2*  *Kcnd2*  *Kcnd3*  *Kcnip2*  *Clock*  *Bmal1*  *Per1*  *Per2*  *Cry1*  *Cry2*  *Dbp*  *Nr1f1*  *Nr1d1*  *Klf15*  *Nfil3*  *Rpl32*  *Tbp*  *Ywhaz*  *ChIP primers*  $\mathrm{RORE}_{1}^{Cacna1c}$  $\mathrm{RORE}_{2}^{Cacna1c}$  RORE^Kcnd2^  $\mathrm{RORE}_{1}^{Kcnd3}$  $\mathrm{RORE}_{2}^{Kcnd3}$ | fwd 5’- CATCATCATCTTCTCCCTCCT -3’  rev 5’- CATCACCGAATTCCAGTCCT -3’  fwd 5’- TTGCAAGAACACTGCAATTGG -3’  rev 5’- GGGCCAGTTTATCAGCTGCTA -3’  fwd 5’- GACCTTGTCACACTGGCAAA -3’  rev 5’- GGCTGCTTGGACTTTCTCTG -3’  fwd 5’- GCCGCAGCACCTAGTCGTT -3’  rev 5’- CACCACGTCGATGATACTCATGA -3’  fwd 5’- CCACCTGCTACACTGCTTAGAA -3’  rev 5’- TCTCCATGCAGTTCTGCTCAAA -3’  fwd 5’- ATGGCTGTATCACGAAGGAG -3’  rev 5’- CCGTCCTTGTTTCTGTCCATC -3’  fwd 5’- CCTATCCTACCTTCGCCACACA -3’  rev 5’- TCCCGTGGAGCAACCTAGAT -3’  fwd 5’- ACATAGGACACCTCGCAGAA -3’  rev 5’- AACCATCGACTTCGTAGCGT -3’  fwd 5’- CTGGCAATGGCAAGGACTC -3’  rev 5’- AGGAGGCTGTAGGCAATGG -3’  fwd 5’- AGCTACACCACCCCTTACAAGCT -3’  rev 5’- GACACGGCAGAAAAAAGATTTCTC -3’  fwd 5’- TTGCCTGTTTCCTGACTCGT -3’  rev 5’- GACAGCCACATCCAACTTCC -3’  fwd 5’- TCGGCTCAACATTGAACGAA -3’  rev 5’- GGGCCACTGGATAGTGCTCT -3’  fwd 5’- AATGACCTTTGAACCTGATCCCGCT -3’  rev 5’- GCTCCAGTACTTCTCATCCTTCTGT -3’  fwd 5’- CGCGGCGTAAAGGATGTATTT -3’  rev 5’- CCACAGATCTTGCATGGAATAATT -3’  fwd 5’- CGTTCGCATCAATCGCAACC -3’  rev 5’- GATGTGGAGTAGGTGAGGTC -3’  fwd 5’- ACAGGCGAGAAGCCCTTT -3’  rev 5’- CATCTGAGCGGGAAAACCT -3’  fwd 5’- GAACTCTGCCTTAGCTGAGGT -3’  rev 5’- ATTCCCGTTTTCTCCGACACG -3’  fwd 5’- GCTGCTGATGTGCAACAAA -3’  rev 5’- GGGATTGGTGACTCTGATGG -3’  fwd 5’- AAAGACCATTGCACTTCGTG -3’  rev 5’- GCTCCTGTGCACACCATTTT -3’  fwd 5’- AGACGGAAGGTGCTGAGAAA -3’  rev 5’- GAAGCATTGGGGATCAAGAA -3’  fwd 5’- GTTTTCCAAAATGGCAGGTCAAGAACC -3’  rev 5’- TGTAGTTTCCTGGTAAGACACCCAGT -3’  fwd 5’- TGGCCTCTGTTCTTGTAAGGTC -3’  rev 5’- CAGCCACACTTAGATTCTTCGC -3’  fwd 5’- GGAGAAAGGTCAAGGCGAGG -3’  rev 5’- TAAGCTGGGAAGAGAGCGAGA -3’  fwd 5’- TGACCTTCTGGCCATGTGTC -3’  rev 5’- GGGAAGATCCCACGGAAGAC -3’  fwd 5’- CTGACCTCAAAGTATGTTTTACCCC -3’  rev 5’- GTTGGGACTTTTCTGAGGGTTG -3’ |

**Supplemental Figure 1**

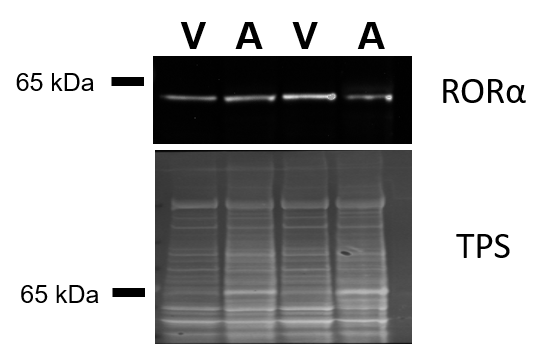

**Supplemental Figure 2.**

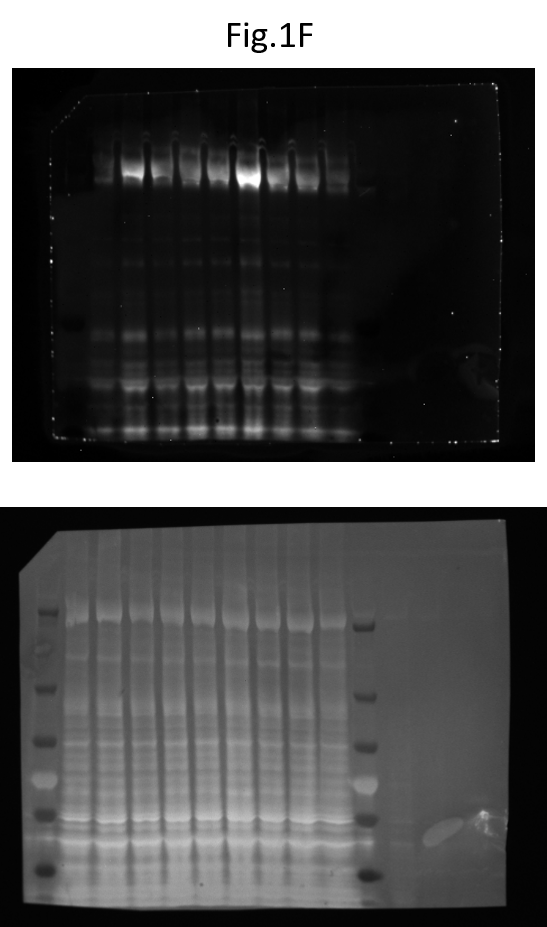

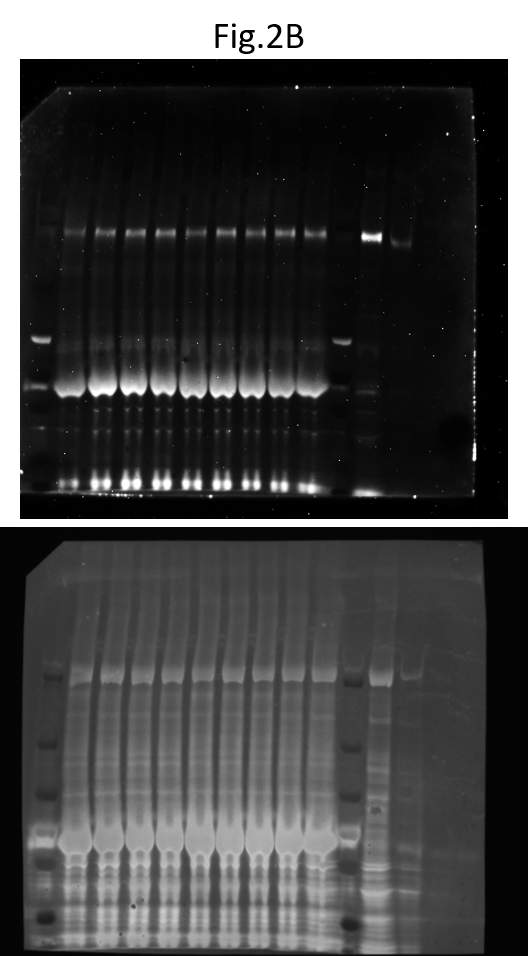

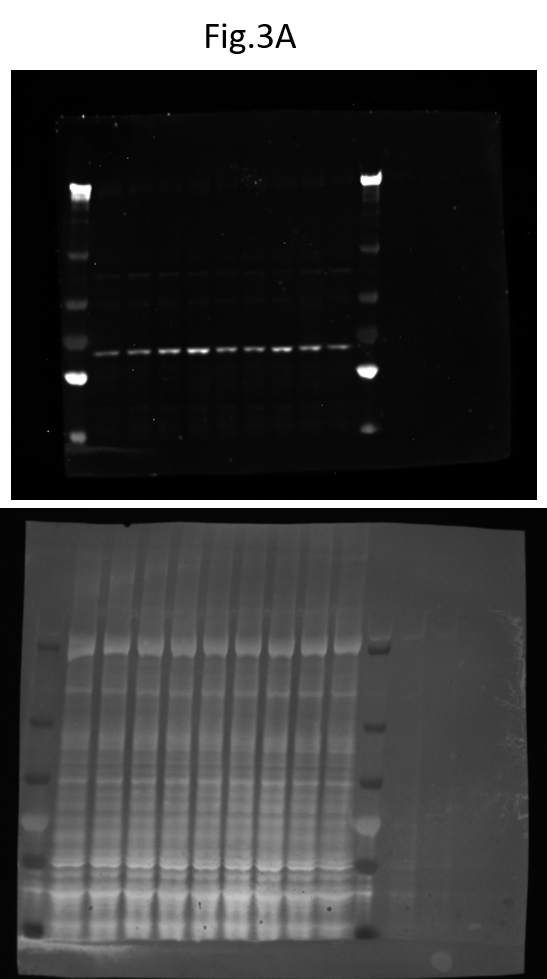

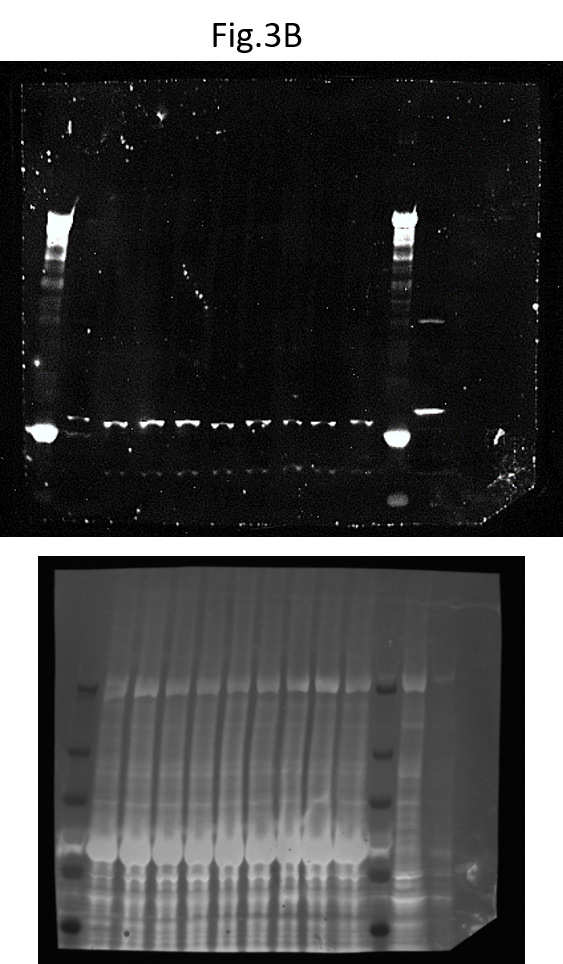

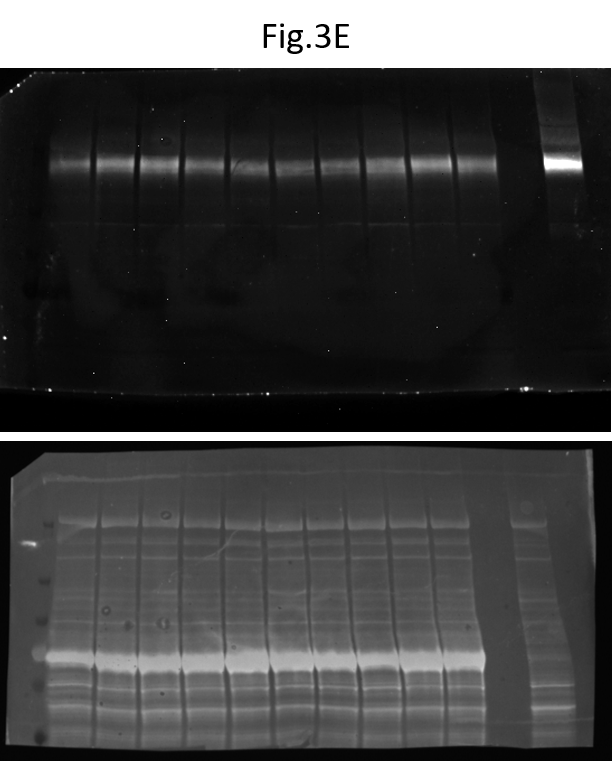

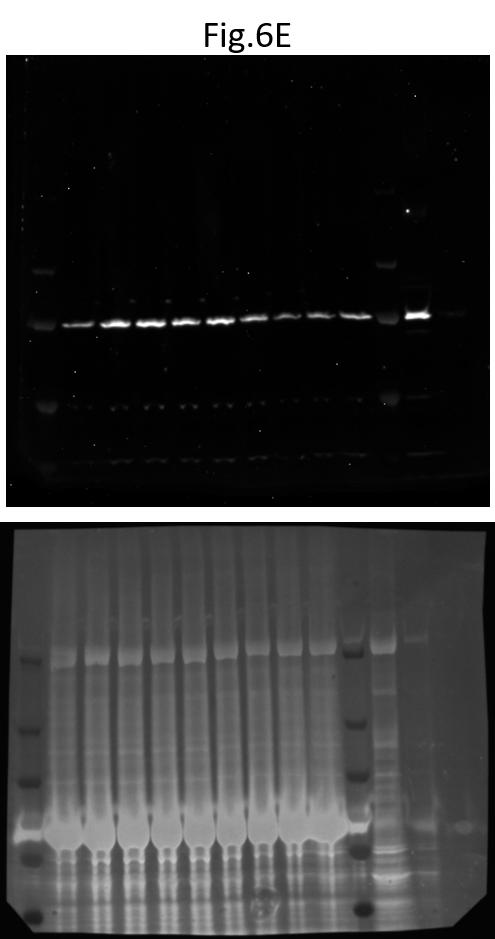

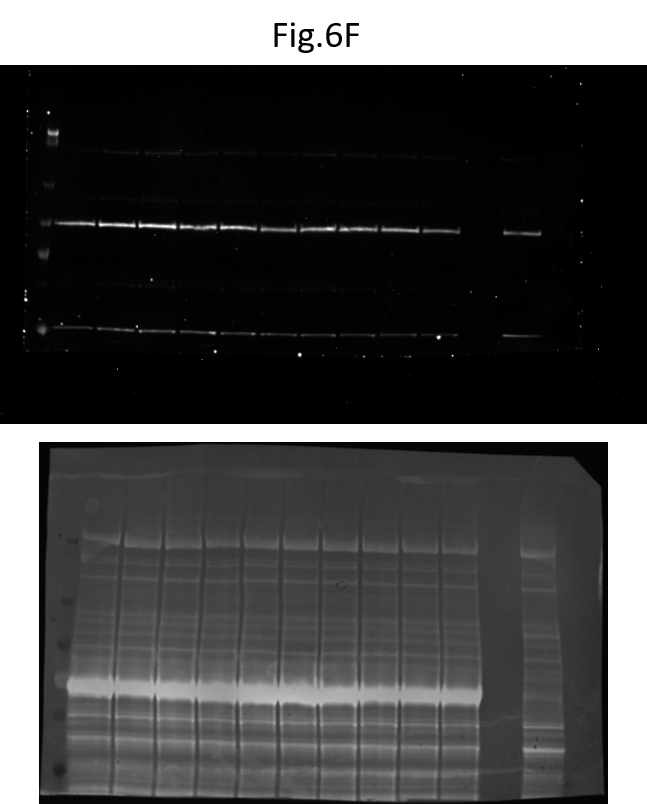
